# The maize pathogen *Ustilago maydis* secretes glycoside hydrolases and carbohydrate oxidases directed towards components of the fungal cell wall

**DOI:** 10.1101/2022.09.16.508353

**Authors:** Jean-Lou Reyre, Sacha Grisel, Mireille Haon, David Navarro, David Ropartz, Sophie Le Gall, Eric Record, Giuliano Sciara, Olivier Tranquet, Jean-Guy Berrin, Bastien Bissaro

## Abstract

Filamentous fungi are keystone microorganisms in the regulation of many processes occurring on Earth, such as plant biomass decay, pathogenesis as well as symbiotic associations. In many of these processes, fungi secrete carbohydrate-active enzymes (CAZymes) to modify and/or degrade carbohydrates. Ten years ago, while evaluating the potential of a secretome from the maize pathogen *Ustilago maydis* to supplement lignocellulolytic cocktails, we noticed it contained many unknown or poorly characterized CAZymes. Here, and after re-annotation of this dataset and detailed phylogenetic analyses, we observed that several CAZymes (including glycoside hydrolases and carbohydrate oxidases) are predicted to act on the fungal cell wall (FCW), notably on β-1,3-glucans. We heterologously produced and biochemically characterized two new CAZymes, called *Um*GH16_1-A and *Um*AA3_2-A. We show that *Um*GH16_1-A displays β-1,3-glucanase activity, with a preference for β-1,3-glucans with short β-1,6 substitutions, and *Um*AA3_2-A is a dehydrogenase catalyzing the oxidation of β-1,3- and β-1,6-gluco-oligosaccharides into the corresponding aldonic acids. Working on model β-1,3-glucans, we show that the linear oligosaccharide products released by *Um*GH16_1-A are further oxidized by *Um*AA3_2-A, bringing to light a putative biocatalytic cascade. Interestingly, analysis of available transcriptomics data indicates that both *Um*GH16_1-A and *Um*AA3_2-A are co-expressed, only during early stages of *U. maydis* infection cycle. Altogether, our results suggest that both enzymes are connected and that additional accessory activities still need to be uncovered to fully understand the biocatalytic cascade at play and its physiological role.

**Importance:** Filamentous fungi play a central regulatory role on Earth, notably in the global carbon cycle. Regardless of their lifestyle, filamentous fungi need to remodel their own cell wall (mostly composed of polysaccharides) to grow and proliferate. To do so, they must secrete a large arsenal of enzymes, most notably carbohydrate-active enzymes (CAZymes). However, research on fungal CAZymes over past decades has mainly focused on finding efficient plant biomass conversion processes while CAZymes directed at the fungus itself have remained little explored. In the present study, using the maize pathogen *Ustilago maydis* as model, we set off to evaluate the prevalence of CAZymes directed towards the fungal cell wall during growth of the fungus on plant biomass and characterized two new CAZymes active on fungal cell wall components. Our results suggest the existence of a biocatalytic cascade that remains to be fully understood.

## INTRODUCTION

Filamentous fungi play a central regulatory role on Earth. Saprophyte fungi, through the decomposition of dead matter, are instrumental in the global carbon cycle while mycorrhizal fungi, by taking in charge the collection and supply of minerals from soils, ensure the survival of most plants via symbiotic associations (1). On the “dark side of the force”, pathogenic fungi, which can cause dramatic crop losses or severe human diseases, also affect ecosystems balance. Significant advances have been achieved in the past decade, notably *via* ambitious genome sequencing programs (2), and post-genomic studies (3, 4), to understand the fungal strategies put in place in these various ecological contexts. This collective corpus of data clearly indicates that during their life cycle filamentous fungi deploy an extraordinary diversity of enzymes, encompassing notably a wide array of carbohydrate-active enzymes (CAZymes) (5–10). In a context largely dominated by the overarching goal of developing efficient plant biomass conversion processes for biorefinery purposes *largo sensu*, the study of the enzymatic arsenal of filamentous fungi has logically focused on enzymes targeting plant components, notably the plant cell wall (PCW). Strikingly, the role of secreted enzymes potentially directed towards the fungus itself has remained under the radar of most studies. Developmental biology of fungi has taught us that to explore their environment, and eventually interact with and/or infect their host, filamentous fungi need to remodel their own cell wall (11). The fungal cell wall (FCW) can also serve as an emergency carbon source, *via* autophagy, in the case of external carbon source shortage (12). Deciphering the potential role of endogenous FCW-targeting enzymes, and their orchestration, is thus of utmost importance. Alike lignocellulose, the FCW is an intricate multi-layer of complex polymers and is depicted today as being composed of an inner layer of chitin, a middle layer of β-1,3-glucans and an outer layer of mannoproteins (13). Some fungal species are reported to also display galactoaminoglycans (14). Several FCW-targeting enzymes secreted by plants as defensive mechanism, notably β-1,3-glucanases and chitinases, have also been reported (15). However, the identity and role of FCW-active CAZymes produced by the fungus itself remains underexplored.

In the present study, we have used as a study model the maize biotrophic pathogen *Ustilago maydis*, also known as corn smut causing major crop yield losses every year (16). *U. maydis* is a rather peculiar filamentous fungus amongst basidiomycetes as it is a dimorphic fungus (i.e. able to switch from yeast to filamentous state). It possesses a rather poor set of lignocellulolytic CAZymes out of a total of 230 CAZymes-encoding genes (www.cazy.org; (17)). Yet, its total secretome produced on maize bran was found to contain 86 proteins, including 23 CAZymes predicted to target the PCW (10). Of note, this latter study on the secretome of *U. maydis* was reported in 2012, i.e. before the extension of the CAZy database with auxiliary activities (AA; (18)) and fine sequence-function understanding of certain GH families, such as GH16 (19). Today, the AA class comprises oxidoreductases that have gained significant importance as they target a wide range of oligo and polysaccharides found in PCW (7) and/or FCW.

In the present study, we have re-analyzed the secretomic data published in 2012 (10) in light of today’s knowledge and identified several enzymes potentially targeting the FCW rather than the PCW. We demonstrate that two of these enzymes, which belong to the GH16 and AA3 CAZy families, are active on β-1,3-glucans or compounds thereof. Our results suggest that both enzymes are most likely involved in a common biocatalytic cascade of importance for the fungus lifestyle.

## RESULTS

### Re-assessment of *U. maydis* secretome on corn bran reveals the presence of putative FCW-active enzymes

The secretome of *U. maydis*, cultivated on corn bran, was first reported in 2012 (10). At that time, the identified top enzymes were arabinoxylan-degrading enzymes (GH10, GH27, GH51, GH62). Here, taking advantage of progresses made in the CAZy field since then, and notably the creation (18) and enrichment of the AA class (20), we set off to re-annotate and evaluate the enzymes deployed by *U. maydis* during the conversion of corn bran. Out of the top 50 proteins, 21 are CAZymes (13 GHs, one expansin, three CE4 and four AAs) (**Fig. 1**). Amongst them, 10 can clearly be predicted as active on PCW (**Table S1**) whereas the roles/targets of the 11 others (two GH5_9s, one GH16, one GH135, three CE4s, three AA3_2s and one AA7) are not so obvious.

**Fig. 1.**
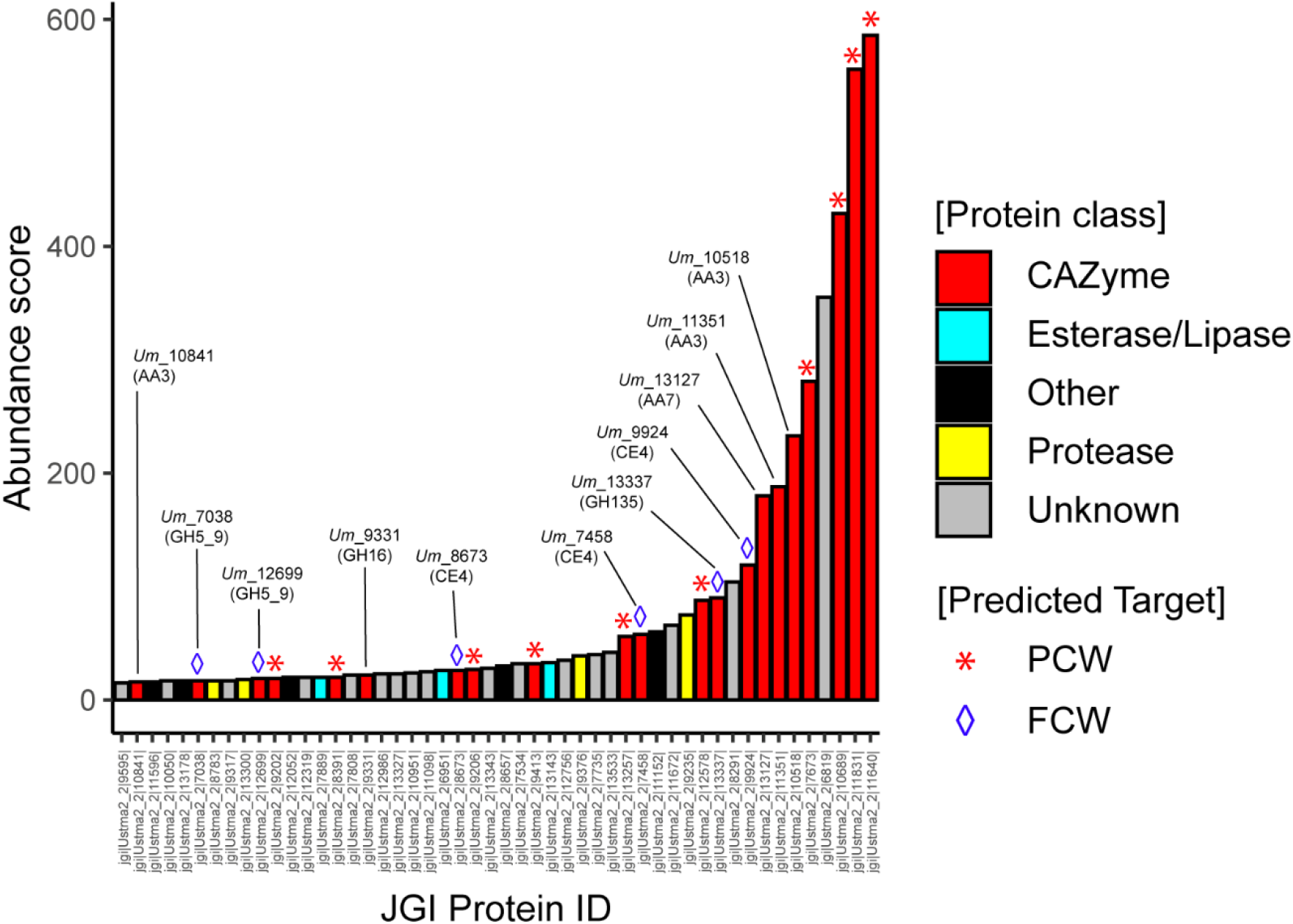
Re-annotation of the Top50 proteins secreted by *U. maydis* when cultivated on corn bran. The enzymes are classified according to their abundance in the secretome (after 7 days growth on maize bran; (10)) and a color code identifies the class of protein (see legend in the figure, “Other” refers to all other types of detected proteins). The protein number that is provided corresponds to the JGI protein ID (*U. maydis* 521 v2.0 strain).

In the present study, our selection of enzymes was guided by: (i) the substrate targeted, and (ii) the likeliness of biological interplay between enzymes. After carrying out phylogenetic analyses, we decided to focus on enzymes predicted to target the main component of the FCW, i.e. the β-1,3-glucans.

Among the putative FCW-active enzymes detected in the secretome (**Table S1**), the three CE4 (*Um*_7458, *Um*_9924 and *Um*_8673; the number corresponds to the JGI protein ID) are directed towards the chitin fraction, as they were biochemically characterized as chitin deacetylases in a recent study by Rizzi et al. (in which they were respectively called *Um*CDA1, *Um*CDA3 and *Um*CDA4) (21). These CE4 were notably shown to be necessary for development and virulence of *U. maydis*. The GH135 (*Um*_13337) is predicted to be active on galactoaminogalactan (GAG), a polysaccharide of the extracellular matrix covering the cell wall.

Regarding the remaining enzyme candidates, to help us in the selection of the most relevant ones for biochemical validation and interplay studies, we searched for hints from biological conditions. To this end, we parsed available data reporting the transcriptomic profiling of the entire life cycle of *U. maydis* on maize (22) (**Fig. S1A**). **Figure S1B** shows the differential transcription along the pathogenic cycle of the genes coding for the 11 CAZymes mentioned above with putative activity on FCW. We observed that one of the AA3_2 (JGI ID 10841/UMAG_03551) and the GH16 (JGI ID 9331/UMAG_02134) are the only ones to display a similar expression profile, and thus a potential interplay: they are expressed at relatively low levels, very early in the cycle (0.5-1 dpi) and are clearly down-regulated during the plant infection cycle.

Phylogenetic analysis of the taxonomically broad GH16 family revealed that, out of 27 sub-families (19), the GH16 *Um*_9331 belongs to the GH16_1 subfamily (**Fig. 2**), and is henceforth called *Um*GH16_1-A (as it is the first GH16 from *U. maydis* to be biochemically characterized). The GH16_1 subfamily is composed of almost exclusively fungal sequences, with the following reported activities: mainly *endo*-β-(1,3)-glucanases (EC 3.2.1.39) *endo-β-* (1,3)/(1,4)-glucanases (EC 3.2.1.6), more seldom hyaluronidase (EC 3.2.1.35) (23) and *exo*-β-(1,3)-glucosyltransferase/elongating β-transglucosylase (EC 2.4.1.–) (24). *Um*GH16_1-A is thus potentially active on β-(1,3)-glucans found in the FCW.

**Fig. 2.**
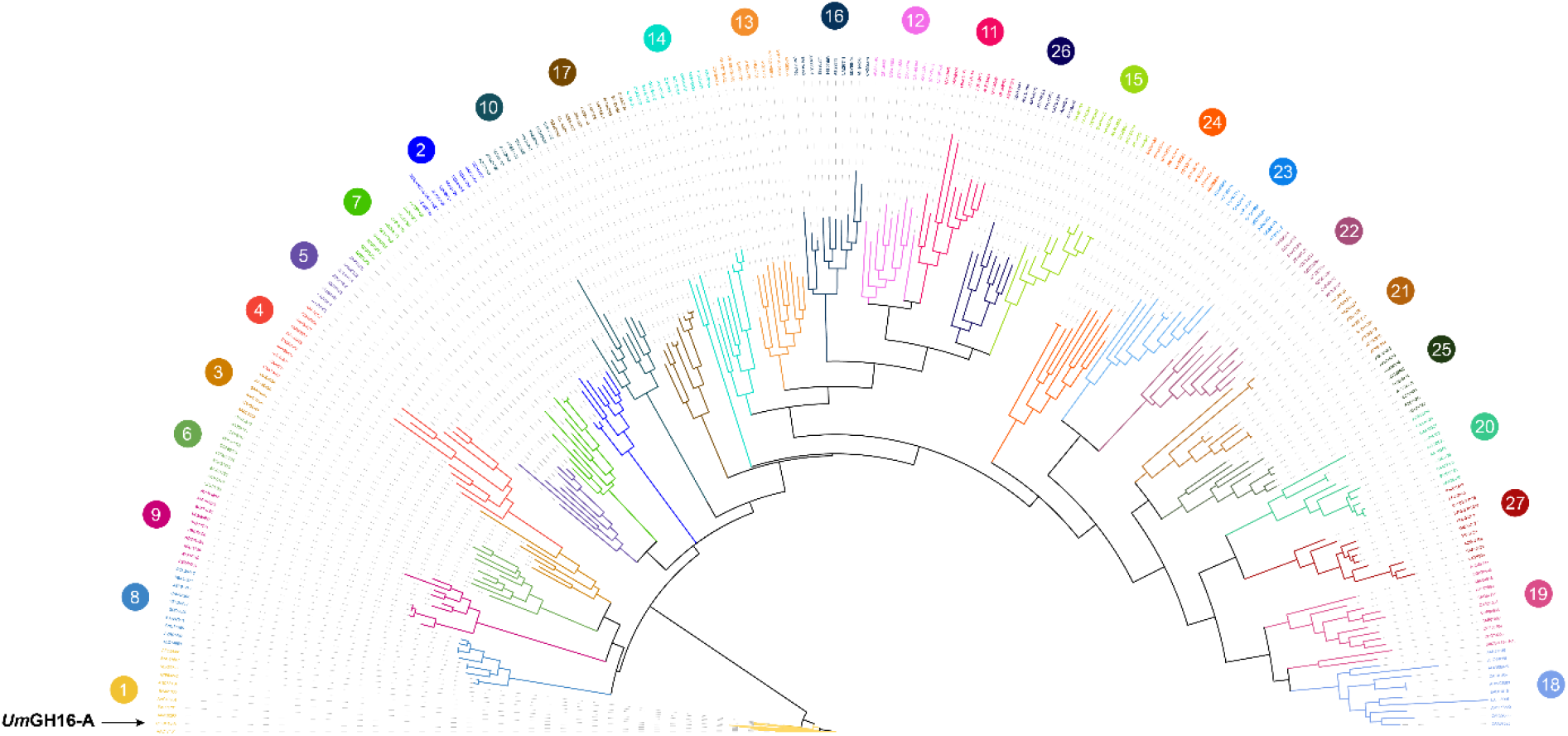
Phylogenetic analysis of GH16 family. Phylogenetic clades, as defined by Viborg et al. (19), are indicated with colored numbers. *Um*GH16_1-A (indicated by a black arrow) falls within the GH16_1 clade. The tree was inferred using RAxML (100 boostraps) on the basis of a MSA made with MAFFT.

The AA3 family is a rather broad family divided into four subfamilies and composed of FAD-dependent oxidases (i.e. main electron acceptor is O2) and dehydrogenases (organic electron acceptor) that oxidize various types of electron donors (25). Phylogenetic analysis revealed that the three AA3s (*Um*_10518, *Um*_10841 and *Um*_11351) detected in the secretome of *U. maydis* all fall within the AA3_2 subfamily (**Fig. 3A**). A closer look at the AA3_2 subfamily (**Fig. 3B**) reveals that *Um*_10518 and *Um*_11351 belong to undefined groups. Interestingly, *Um*_10841 (henceforth called *Um*AA3_2-A) clusters together with the AA3_2 from the white-rot fungus *Pycnoporus cinnabarinus* hitherto called *Pc*GDH (26), and recently renamed oligosaccharide dehydrogenase (ODH) after it was shown to be active on laminaribiose (G3G; Glc-β-1,3-Glc) (27). Of note, ODH appeared to be much more active on G3G than on the initially reported substrate, *viz*. glucose. This recent finding highlights that the phylogenetic functional annotation and biological role of AA3_2s is far from being firmly established and that *Pc*ODH and *Um*AA3_2-A may form a new, intermediate clade in between GOX and GDH activities. While this analysis suggests that *Um*AA3_2-A could be a candidate for the oxidation of β-1,3-glucans components, its rather diverging sequence (46% sequence identity with *Pc*ODH) called for biochemical investigations.

**Fig. 3.**
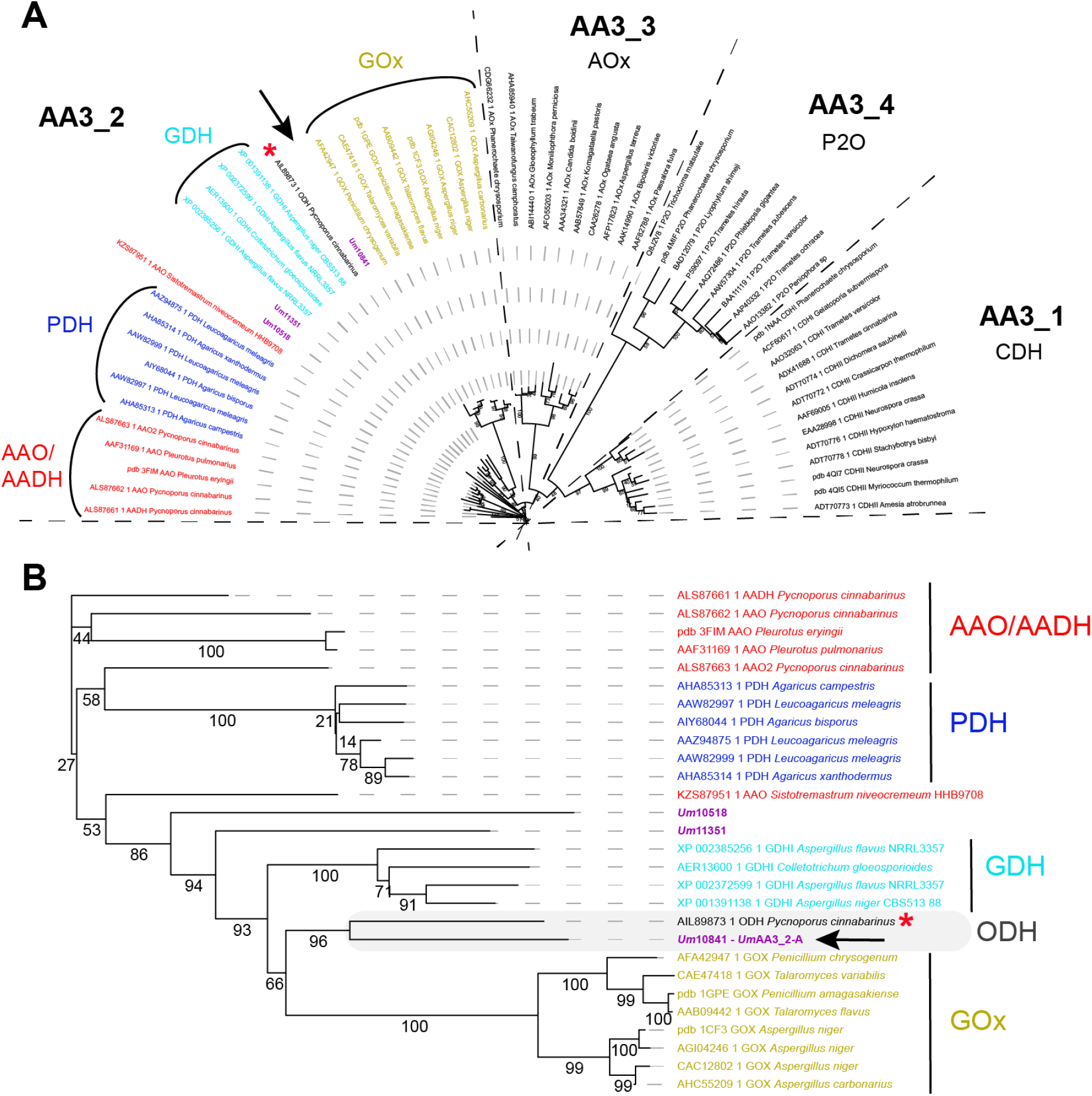
Phylogenetic analysis of the AA3 family (A) and zoom-in view on the AA3_2 subfamily (B). The AA3s identified in the secretome of *U. maydis* are shown in purple. The new oligosaccharide dehydrogenase clade, including *Um*AA3_2-A (indicated by a black arrow), characterized in the present study, and the *Pc*GDH (red asterisk), is framed in grey. The tree was inferred using PhyML (bootstrap values, as percent, are shown on the branches).

### *Um*GH16_1-A is a β-1,3-glucanase with preference for β-1,3-glucans branched with short β-1,6 substitutions

*Um*GH16_1-A was heterologously expressed in *Pichia pastoris* and purified to homogeneity (**Fig. S2**) but the protein yield was very low (0.175 mg/L of culture), preventing extensive characterization. Sequence and structure comparisons between *Um*GH16_1-A (Alpha-fold2 model; (28)) and its closest structural homologue (RMSD of 0.473 Å and sequence identity of 38%), *viz*. the GH16_1 from *Phanerochaete chrysosporium* (*Pc*GH16_1; also called Lam16A; PDB id 2W52; (29)), indicated the presence in *Um*GH16_1-A of an extra 57 amino acid C-terminal extension with no predictable canonical fold (**Fig. S3**). A protein blast search on ncbi against the nr database showed the occurrence of orthologs of *Um*GH16_1-A bearing similar C-terminal extensions in a broad range of *Ustilaginomycotina* fungi (data not shown). We hypothesized that this extension could pose heterologous production issues and found that, indeed, upon its deletion, the production of the catalytic domain (cd) of *Um*GH16_1-A, henceforth called *Um*GH16_1-A_cd, was increased by ca. 40-fold (ca. 7 mg/L of culture).

Screening of the substrate specificity of *Um*GH16_1-A_cd showed the release of oligosaccharides from different β-1,3-glucans, with the highest amounts of products detected for laminarin, followed by yeast β-glucan and then pachyman (**Fig. 4A&B**). In order to further understand this substrate preference, we carried out linkage analysis of those three substrates (**Fig. 4C, Fig. S4A&B)**. We confirm that Pachyman is a linear β-1,3-glucan and show that the structure of laminarin and yeast β-glucans is somewhat different from the suppliers’ descriptions. Indeed, laminarin appears to be a linear β-1,3-glucan with low frequency (ca. 3%) of single glucose units branched in β-1,6 (**Fig. 4C**). Interestingly, yeast β-glucan appears to have a similarly low substitution frequency, but longer branches (on average four β-1,6-linked glucose units on each branch).

**Fig. 4.**
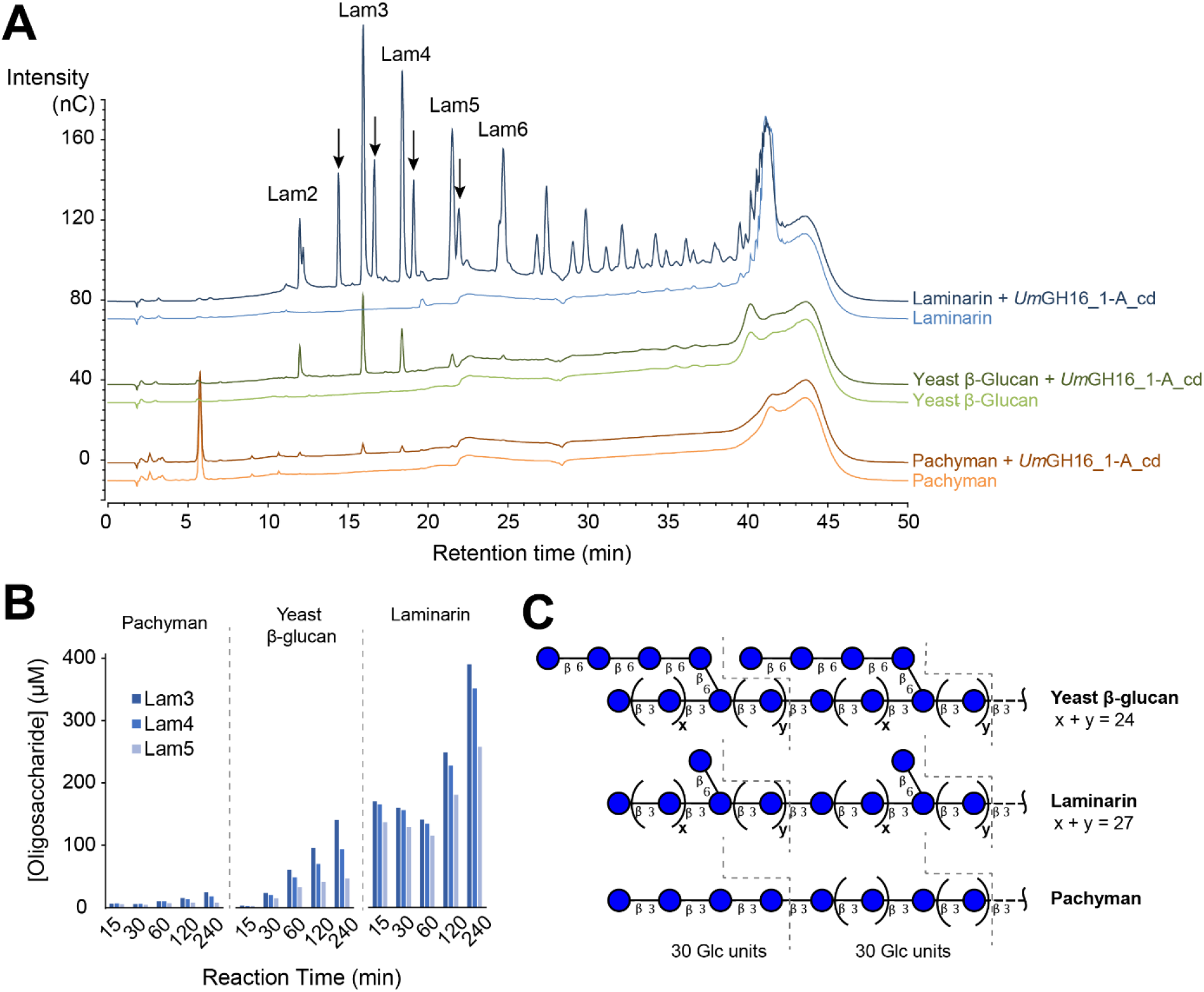
Activity of *Um*GH16_1-A_cd on β-1,3 glucans. **(A)** The graphs show HPAEC-PAD chromatograms of reaction products released from laminarin, yeast β-glucan and pachyman (10 mg.mL^-1^ final concentration) by *Um*GH16_1-A_cd (10 nM). Black arrows indicate reduced β-1,3-gluco-oligosaccharides (see **Fig. S6**). All reactions were incubated during 4 h, in citrate phosphate buffer (50 mM, pH 5.5), in a thermomixer (30 °C, 1,000 rpm). All experiments were carried out in triplicate but for the sake of clarity, only one replicate is shown. See **Fig. S7** for additional control experiments. **(B)** Time-course release of Lam3-Lam5 oligosaccharides from laminarin, yeast β-glucan and pachyman (same reaction conditions as in panel A; n = 1). **(C)** Proposed chemical structure of the three tested polymers on the basis of carbohydrate linkage analysis (see **Fig. S4** for more details). «β3 and «β6» represents β-(1,3) and β-(1,6) linkages.

We underscore that HPAEC-PAD and LC-MS analyses did not show the release of β-1,6/1,3-gluco-oligosaccharides. This is in contrast with the product profile of its ortholog *Pc*GH16_1 acting on laminarin, for which the release of G6G3G3G has been shown by NMR (30). Yet, the presence of short β-1,6 branches on the β-1,3-glucan main chain appears to increase significantly the activity of *Um*GH16_1-A_cd (**Fig. 4A-B)**. We propose that the presence of those substitutions may either help the enzyme to bind to the targeted β-1,3-chain and/or alter the physicochemical properties of the polymer improving reactivity with the enzyme.

Furthermore, one can observe with both HPAEC-PAD (**Fig. 4A**) and MALDI-ToF MS (**Fig. S5**) analyses the release from laminarin by *Um*GH16_1-A_cd of a series of secondary peaks adjacent to the Lam series. LC-MS analysis of these peaks showed that they correspond to C1-reduced cello-oligosaccharides, already present in the initial laminarin suspension (**Fig. S6**). This modification most probably occurred during laminarin extraction/preparation by the supplier.

Control experiments showed no (for DP2-DP4) or extremely low (for DP5-DP6) activity on β-1,3-gluco-oligosaccharides (**Fig. S7A&B**). This observation suggests that the enzyme requires more than six carbohydrate units to be active. Furthermore, the concomitant release from laminarin of oligosaccharides with both low and high DP by *Um*GH16_1-A_cd, even at very early time points (**Fig. S8**), suggests that the enzyme would act with both *exo* and *endo* modes. This question would deserve further investigations. Additional control experiments showed that no activity could be detected on any of the tested polysaccharides with β-1,4 linkages (cellulose and α-chitin) or mixed β-1,3/1,4 linkages (lichenan) (**Fig. S7C**).

Overall, we conclude that *Um*GH16_1-A_cd is an β-1,3-glucanase with a marked preference for β-1,3-glucans substituted with single β-1,6-branched glucose units.

### *UmAA3_2-A* is a dehydrogenase active on β-1,3 and β-1,6-gluco-oligosaccharides

*Um*AA3_2-A was successfully heterologously produced in *P. pastoris* and purified to homogeneity (ca. 5 mg/L of culture). We underscore that SDS-PAGE analysis was not trivial, most likely due to excessive protein stability under denaturating conditions (main band with apparent MW ~ 50 kDa), proteolysis (band at ~ 25 kDa) and oligomerization mediated by inter-molecular disulfide-bonds (**Fig. S9A**). During the preparation of this manuscript, Wijayanti et al. reported the production and preliminary characterization of several AA3_2s, including *Um*AA3_2-A (called there *Um*GDHIII; (31)), for which they observed the same atypical, cleavage and polymerization pattern under SDS-PAGE denaturating conditions. We carried out size exclusion chromatography and showed a unique, monodisperse peak, corresponding to an estimated monomeric size of 48 kDa (**Fig. S10B**).

*Um*AA3_2-A substrate specificity was assessed (**Fig. 5A**) by measuring dehydrogenase activity (with DCIP as an electron acceptor) on various (oligo)saccharides **(Fig. S10)**. This analysis revealed that β-d-Glc*p*-(1,6)-d-Glc (G6G, also called gentiobiose), followed by β-d-Glc*p*-(1,3)-d-Glc (G3G; laminaribiose) and G3G3G (laminaritriose) were the preferred substrates. These results are in good agreement with those reported by Cerutti et al. (27) and Wijayanti et al. (31). As in those two works, no activity on either cellobiose (G4G), cellotriose (G4G4G) or the trisaccharide G3G4G could be measured, whereas some activity was retained on the mixed trisaccharide G4G3G containing a β-(1,3) glycosidic bond between the reducing end glucose unit and the adjacent unit (**Fig. 5A)**. We determined an optimum pH of 5.5 for the dehydrogenase activity, on glucose (**Fig. S11A**), G3G (**Fig. S11B**) and G6G (**Fig. S11C**). Of note, we also probed the ability of *Um*AA3_2-A to reduce O2 in the absence of organic electron acceptor by measuring the production of H2O2 using either G3G, G6G or glucose as electron donor (**Fig. 5B**). No oxidase activity could be detected, confirming thereby the strict dehydrogenase nature of the enzyme as previously observed in hereinbefore mentioned studies (26, 27, 31).

**Fig. 5.**
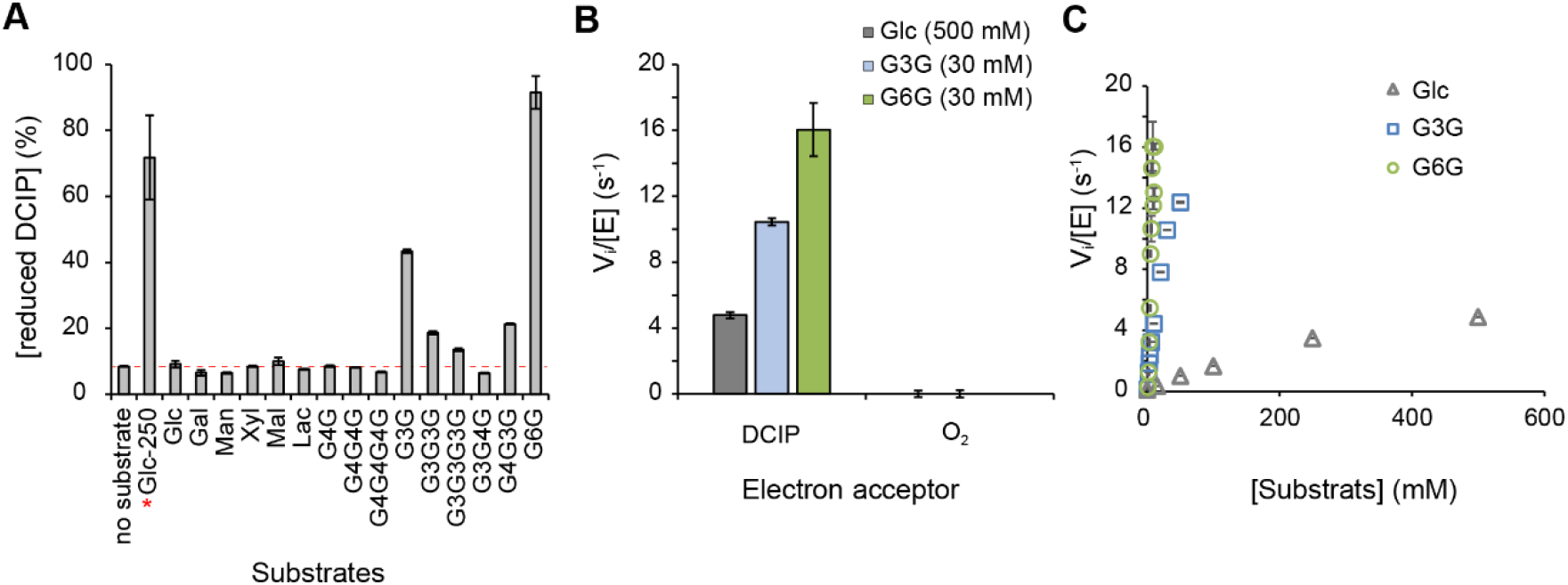
Activity of *Um*AA3_2-A. **(A)** Substrate specificity screening monitored as the reduction of DCIP (400 μM) by *Um*AA3_2-A (14 nM) in the presence of various substrates (2.5 mM for all, 250 mM when marked with a red star) after 3h incubation (see **Fig. S10** for substrate nomenclature). **(B)** Dehydrogenase *vs* oxidase activity measured respectively as the reduction of DCIP (400 μM) *vs* O_2_ (250 μM) by *Um*AA3_2-A (110 nM) in the presence of Glucose (500 mM) or G3G (30 mM). **(C)** [Glucose] and [G3G]-dependency of *Um*AA3_2 initial rate. All reactions were carried out in citrate-phosphate buffer (50 mM, pH 5.5), at 30 °C. Data are presented as average values (n = 3, independent biological replicates) and error bars show s.d.

We then determined the dehydrogenase kinetic parameters of *Um*AA3_2-A, at optimum pH, for glucose, G3G and G6G (**Fig. 5C**). For these three substrates, substrate saturation could hardly be reached within a reasonable concentration range. Yet, determination of the initial slopes on the Vi = f(S) plot allowed us to approximate the catalytic efficiencies, yielding values of *k*cat/*K*M of 697, 636 and 18 M^-1^.s^-1^ for G6G, G3G and glucose, respectively (see **Table S2** for the full set of approximate kinetic parameters).

Here, in addition to the commonly used DCIP-based assay, we used LC and MS methods to characterize the product profile of this enzyme. Using mass spectrometry, we first verified that the reaction catalyzed by *Um*AA3_2-A on G3G, G3G3G and G6G yielded oxidized species, as shown by the presence of simple and double sodium adducts of M+16 species (**Fig. S12**). To establish whether these species are geminal-diols (i.e. oxidized on non-reducing end carbon) or aldonic acids (i.e. oxidized on the C1 carbon of substrate reducing end) we carried out UPLC-MS using positive and negative ionization mode (**Fig. S13-S15**). For conversion reactions of G3G (**Fig. S13**), G3G3G (**Fig. S14**) and G6G (**Fig. S15**), one oxidized species was observed, in negative mode only, which is indicative of the formation of the corresponding aldonic acid.

Altogether, these results are consistent with a two-electron oxidation of the oligosaccharide at the C1 carbon, yielding a lactone, which is known to undergo a spontaneous hydrolysis leading *in fine* to aldonic acids.

For comparison purposes, Cerutti et al. reported for *Pc*ODH *k*cat/*K*M values of 777 M^-1^.s^-1^ and 47 M^-1^.s^-1^for G3G and glucose, respectively (27). Wijayanti at al. reported similar kinetic parameters for *Um*AA3_2-A as those we present here (see **Table S2**). For both *Pc*ODH and *Um*AA3_2-A, the presence of a β-1,3 linkage between the reducing and first non-reducing D-Glc units is thus clearly crucial for the activity.

To gain insight into the structure-function relationship underlying *Um*AA3_2-A mode of action, we compared a homology model (generated using AlphaFold) to the X-ray structure of the *Pc*ODH-G3G complex (PDB id: 6XUV; (27)) (**Fig. S16**). This analysis shows very similar structure and active site architecture between *Um*AA3_2-A and *Pc*ODH, with a wider active site entrance than the one observed for *An*GOX and *Af*GDH that accommodate monosaccharides. In particular, Y64, F416 and W430, as well as F421 from the flexible “substrate binding loop” described for *Pc*ODH, are held in optimal position to bind the reducing and non-reducing end, respectively, of G3G, by CH-π interactions. This observation correlates with better dehydrogenase activity detected on β-1,3-oligosaccharides than on glucose. In line with this, out of three residues involved in hydrogen bonding to glucose hydroxyl groups in *Af*GDH and *AnGOX*, and lacking in *Pc*ODH, only one residue (Asp446) is conserved in *Um*AA3_2-A (**Fig. S17**). Remarkably, this residue interacts with glucose O4 hydroxyl in *Af*GDH (Glu435), and seems to be strictly conserved in type I GDH and in GOx enzymes (Asp446 in *An*GOx), as well as in most ODH and ODH-like enzymes, with a few exception, such as *Pc*ODH (Val428) (27). This comparison also revealed the presence of an additional loop (residues 173-192) in *Um*AA3_2-A (**Fig. S17**), that seems conserved in ODH-like proteins as previously described (27). The role of these structural differences in potential biocatalytic differences and biological functions remains to be investigated.

### *Um*GH16_1-A and *Um*AA3_2-A interplay on fungal β-1,3 glucans

As shown above, *Um*GH16_1-A and *Um*AA3_2-A are active on β-1,3/β-1,6-glucans and oligosaccharides thereof, respectively. Therefore, we set off to investigate the interplay between both enzymes. Using a fraction of *U. maydis* fungal cell wall (*Um*FCW) enriched in β-1,6/β-1,3-glucans, we could detect the release of β-1,3-gluco-oligosaccharides by the versatile, commercial *Tsp*GH16_3 but could not detect any activity when using *Um*GH16_1-A_cd, despite several attempts (**Fig. S18**). We suspect that the nature and branch length of substitutions present in *Um*FCW β-glucans hamper *Um*GH16_1-A-cd activity, underlining the necessity to finely characterize the FCW fraction. Working with a better characterized glucan polymer (i.e. laminarin), we demonstrated that the *in vitro* combination of *Um*GH16_1-A-cd and *Um*AA3_2-A led to a functional biocatalytic cascade where β-1,3-gluco-oligosaccharides released by *Um*GH16_1-A-cd (DP2 to > DP6) were further oxidized by *Um*AA3_2-A (**Figs. 6 & 7**).

**Fig. 6.**
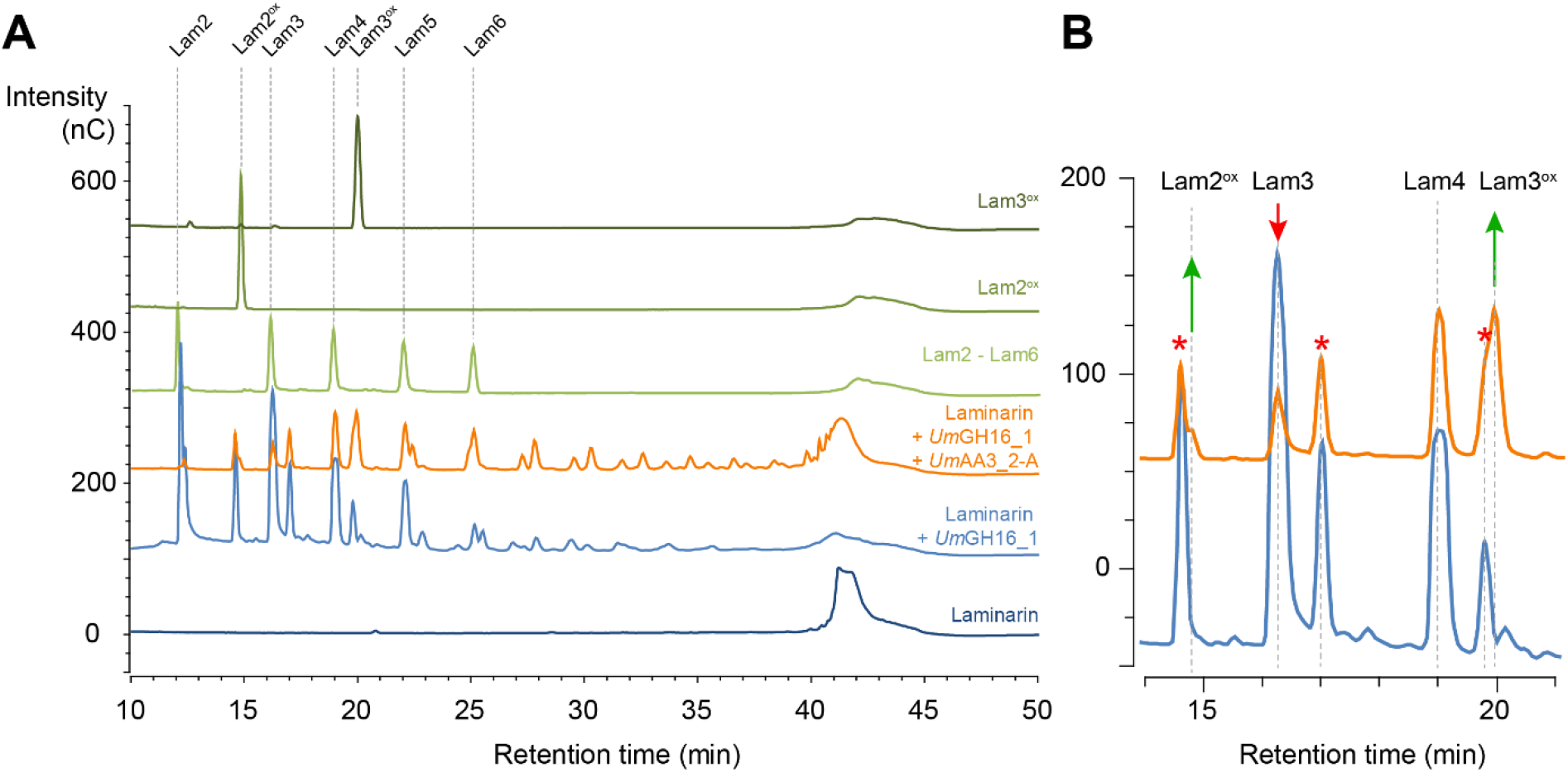
Combined action of *Um*GH16_1-A and *UmAA3_2-A* on laminarin. **(A)** Full HPAEC-PAD chromatogram and **(B)** zoom-in view on the 11-21 min region comparing products released from laminarin by *Um*GH16_1 alone (blue line) or in combination with *Um*AA3_2-A (orange line). The red stars indicate peaks of reduced oligosaccharides already present in the laminarin (see main text).

**Fig. 7.**
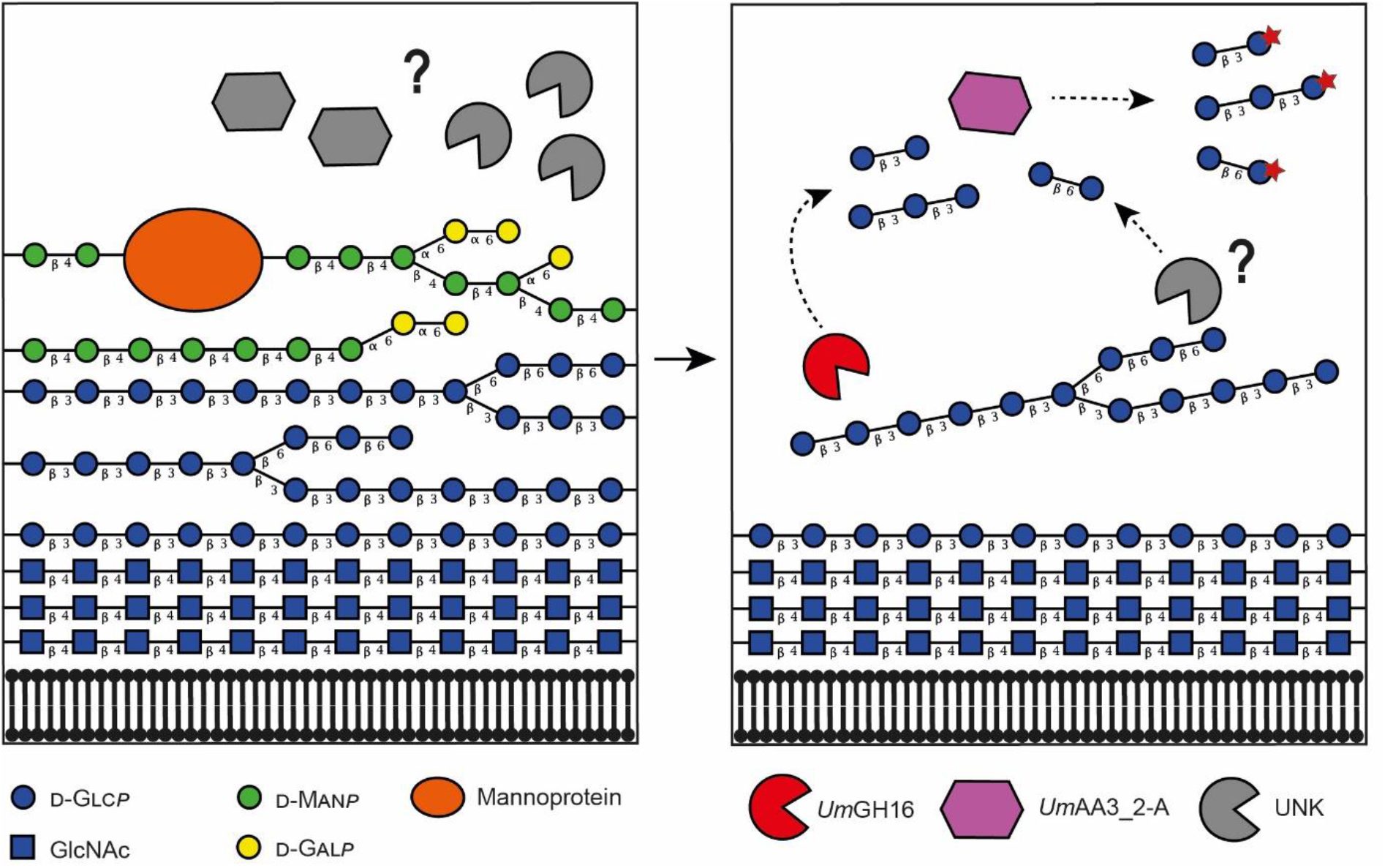
Proposed reaction scheme illustrating the combined action of *Um*GH16_1-A and *UmAA3_2-A* on FCW. Putative enzymatic activities secreted by *Ustilago maydis* to target its own cell wall. The legend key and the glycosidic linkage between each carbohydrate unit are indicated in the figure (for instance, “β 4” indicates a β-1,4 linkage). In the left-hand side panel, hypothetical enzymatic activities (in grey) degrade the galactomannan and mannoproteins, allowing access to the lower layer of β-1,3/β-1,6-glucans. In the right-hand side panel, the uncovered glucans can act as potential substrate for *Um*GH16_1-A (shown in red) and other hypothetical hydrolytic activities (in grey), releasing β-1,3 and β-1,6-oligosaccharides oxidizable by *Um*AA3_2-A (in purple).

To get further insights into the relevance of this potential interplay, we tested two hypotheses. In our first hypothesis, we tested whether product inhibition of the GH16 enzyme by its oligosaccharide products could be alleviated upon their oxidation by the dehydrogenase. A similar scenario has been observed for the cellobiose hydrolase/cellobiose dehydrogenase pair, where cellobiose released from cellulose by the cellobiose hydrolase is no longer an inhibitor for the latter upon oxidation by the cellobiose dehydrogenase (32). However, here, *Um*GH16_1-A_cd was neither inhibited by G3G (**Fig. S19A**), nor by G6G (**Fig. S19B**). Conversely, in our second hypothesis, oxidized oligosaccharides, generated by *Um*AA3_2, could be inhibitors of *Um*GH16-A. The addition of G3G^ox^ or G6G^ox^ to a reaction of *Um*GH16_1-A_cd on laminarin did not show any significant inhibitory effect (**Fig. S20**). Thus, our results rule out any product-based regulatory interplay between both enzymes.

## DISCUSSION

In the present study, by re-analyzing previously published data (10) in the light of today’s knowledge, we have revealed that the secretome of the plant pathogen *U. maydis* grown on corn bran contains a significant fraction of CAZymes predicted to be active on the FCW, including several hydrolases and carbohydrate oxidases that may act in concert. On the basis of phylogenetic analyses and after interrogating published transcriptomic studies, we have selected and biochemically characterized *Um*GH16_1-A and *Um*AA3_2-A, which proved to be active on β-1,3/1,6-glucans and oligosaccharides thereof, respectively.

Together with previously published work (27, 31), we show that both *Pc*ODH and *Um*AA3_2-A appear to form an evolutionarily distinct clade between AA3_2 GOx and GDH, associated to a new substrate specificity. Enzyme kinetics tell us that G6G and G3G are > one order of magnitude better substrates than glucose. Yet, the measured rates still remain low compared to other AA3 dehydrogenases, indicating that the biologically relevant substrate may be more complex, potentially harboring some ramifications. Beyond the structure of the natural substrate, there are also open questions regarding the fate of the electrons extracted from G3G or G6G by *Um*AA3_2. Analogous enzymatic systems active on β-(1,4)-glucans (cellulose and cello-oligosaccharides) have shown that the extracted reducing power could feed downstream enzymatic activities such as LPMOs (33–35). Provided it exists, a similar cascade remains to be found for β-(1,3)-glucans.

Furthermore, while apparent activity on β-1,3/1,6 glucans led us to focus on FCW, it is worth mentioning that β-1,3 glucans can also be found in the cell walls of cereals grasses (including maize) but in the form of mixed-linkage (1→3),(1→4)-β-d-glucans, mostly concentrated in the endosperm (36, 37). Although we cannot rule out these PCW components as potential target for the enzymes studied herein, several facts rather support the hypothesis of FCW-directed activities: (i) *Um*GH16_1-A being active on β-1,3/1,6-glucans but not on lichenan (a mixed β-1,3/1,4 linkages polymer), and *Um*AA3_2-A being most active on G6G, together with (ii) β-1,3/1,6-glucans being mainly present in FCWs (as well as in some seaweeds and bacteria) (38), and virtually absent from plants, and (iii) their co-expression during early stages of *U. maydis* infection cycle and repression at later stages during plant infection cycle by *U. maydis*. While additional accessory activities still need to be uncovered to fully understand the biocatalytic cascade at play (**Fig. 7**), we believe *Um*GH16_1-A and *Um*AA3_2-A could play a role in FCW remodelling, during which released fungal oligosaccharides, known to act as elicitors of plant immunity (39), may be oxidized to evade the host immune response. Indeed, the fate and role of FCW/PCW-derived oxidized products release by fungal oxidative enzymes is an emerging matter of utmost importance (34, 40–43).

## CONCLUSIONS

Recent -omics studies and biochemical characterization have enriched our knowledge over the plethora of activities that constitute the fungal enzymatic arsenal. The various questions that emerge from our study underscore the need for a deeper integration of enzymology, cellular biology and microbial ecology to better understand the genuine activities, biological role, and potential biotechnological interest of CAZymes, and most notably of oligosaccharide oxidases. We believe that the diversity and roles of FCW-active enzymes only starts to unfold, promising important discoveries to be made in the coming years.

## MATERIAL AND METHODS

### Materials

Most chemicals were purchased from Sigma-Aldrich. Oligosaccharides substrates and polysaccharides (Yeast β-glucan, reference P-BGYST, batch number: 180808a / pachyman, reference P-PACHY, batch number: 10301/Lichenan reference P-LICHN, batch number: 70901b), as well as the *endo*-1,3-β-d-glucanase *Tsp*GH16 (reference E-LAMSE) were purchased from Megazyme (Wicklow, Ireland). Laminarin was purchased from Merk (reference L9634).

### Enzymes cloning, production and purification

The gene encoding *Um*AA3_2-A (Uniprot ID A0A0D1DW37, Gene ID UMAG_03551) was PCR amplified from the genome of *Ustilago maydis* BRFM 1093 strain, with the following primers containing EcoRI and XbaI restriction sites (underlined):

Forward: <u>GAATTC</u>GCCATCGTCACAGATG
Reverse: <u>TCTAGACC</u>CCTGGCGAGAATGGTGT

The amplicon and TOPO vector were subsequently used to co-transform *E. coli* DH5α competent cells according to the TOPO^®^ Cloning reaction protocol (Invitrogen). Positive transformants were selected on LB-agar-ampicillin (50 μg.mL^-1^). Plasmidic DNA was extracted, purified and the expected size was verified by agarose electrophoresis. Then, the pPICZαA vector and TOPO-*Um*AA3_2-A vectors were digested with EcoRI and XbaI, gel-purified and a ligation of linearized pPICZαA and *Um*AA3_2-A insert was carried out. The ligation product was then transformed in *E.coli* DH5α for plasmid production. After plasmid extraction the final construct pPICZαA-*Um*AA3_2-A was sequenced before transformation in *P. pastoris*. The intron-free sequence of the gene coding for *Um*GH16_1-A (Uniprot ID A0A0D1E047, Gene ID UMAG_02134) was synthesized after codon optimization for expression in *P. pastoris* and inserted into a modified pPICZαC vector using *XhoI*\* and *NotI* restriction sites in frame with the α secretion factor at N-terminus (i.e. without native signal peptide) and with a (His)_6_-tag at the C-terminus (without *c*-myc epitope) (Genewiz, Leipzig, Germany). For both enzymes, transformation of competent *P. pastoris* X33 was performed by electroporation with PmeI-linearized pPICZαC recombinant plasmids. *P. pastoris* strain X33 and the pPICZαC vector are components of the *P. pastoris* Easy Select Expression System (Invitrogen), all media and protocols are described in the manufacturer’s manual (Invitrogen). Zeocin-resistant *P. pastoris* transformants were screened for protein production as described by Haon et al. (44). The best-producing transformants were grown in 2 L flasks. The proteins of interest were expressed and secreted upon methanol induction and purified from the supernatant by IMAC, according to a previously described protocol (45).

Enzyme concentrations were determined by measuring UV absorbance at 280 nm using a Nanodrop ND-200 spectrophotometer (Thermo Fisher Scientific, Massachusetts, USA) and extinction coefficient determined with ProtParam (Expasy) for *Um*GH16_1-A (ε^280^ = 83,350 M^-1^.cm^-1^) and *Um*AA3_2-A (ε^280^ = 85,830 M^-1^.cm^-1^).

### Phylogenetic Analyses

To build the phylogenetic tree of AA3s, we used 57 sequences of experimentally characterized enzymes of fungal origin (ascomycetes and basidiomycetes) belonging to the four different subfamilies (i.e. AA3_1 to AA3_4), together with the three AA3_2s from *U. maydis* secretome (JGI ID 10518, 10841 and 11351). For the GH16s tree, 264 sequences (including *Um*GH16-A) representing the 27 subfamilies described in the work of Viborg et al., were provided by the CAZy team (AFMB, Marseille), (19). Of note, the variable C-terminal regions of the GH16s sequences were cut using BioEdit (46) in order to keep the catalytic domain only. Both AA3 and GH16 sequences batches were aligned using MAFFT-DASH (L-INS-i method) (47), which include structural data input. The resulting multiple sequence alignments were used to infer the phylogenetic trees via the MAFFT online platform for AA3s and the RAxML software for GH16s. A neighbour-joining method (NJ, on the basis of conserved sites) or a Maximum Likelihood method (ML) was used for AA3s and GH16s, respectively. In both cases, the Whelan and Goldman (WAG) amino acid substitution model was selected (48). Branch support was calculated by 500 (for the AA3s tree, values displayed in percent on the tree) or 100 (for the GH16s tree) bootstrap repetitions. The trees were visualized in iTOL (49) and edited in Illustrator^®^.

### Fungal Cell Wall extraction

The *U. maydis* strain 521, which was provided by the CIRM-CF collection (strain BRFM1093) (50) was grown in 100 mL of Yeast Extract (10 g.L^-1^) / BactoPeptone (20 g.L^-1^) / Dextrose (20 g.L^-1^) (YPD medium) for 48 h at 28 °C in 250 mL-baffled Erlenmeyer flask under orbital agitation (150 rpm). Cells were then harvested and washed once in H2O by centrifugation (1,500 *g*, 10 min), counted and stored at 10^7^ cells/mL in 20% glycerol at −80 °C as a working cell bank for long term preservation. In order to produce material for sequential extraction twenty 250 mL-baffled Erlenmeyer flasks containing 100 mL of YPD medium were inoculated at 10^5^ cells/mL with *U. maydis* cells from the frozen cell bank and incubated for 24h at 28 °C under orbital agitation (150 rpm). Cells were then harvested and washed three times with H2O by centrifugation cycles (8,000 *g*, 4 °C, 20 min). The washed cell pellet was lyophilized (ca. 7 g). Five grams of this material was resupended in 500 mL H2O, homogenized using utra-turax (2 min, 13,500 rpm) and boiled for 4 h. After a centrifugation (8000 *g*, 20 min), the supernatant was filtered on 0.7 μm glass microfibers and stored at 4 °C. The pellet was resupendend in 500 mL of 1.25 M NaOH solution for 4 h at 60 °C. After centrifugation (8000 *g*, 20 min), the supernatant was filtered on 0.7 μm glass microfibers and stored at 4 °C. Polysaccharides extracted in H2O and NaOH were futher precipited in 50% ethanol at 4 °C for 16 h under stirring. Precipitated polysaccharides were washed five times with 50 mL of 50% ethanol and lyophilized. Alkali insoluble material was washed in H2O until pH reached 7 and kept in suspension to enable pipetting.

### Dehydrogenase activity assay

The dehydrogenase activity was monitored by measuring spectrophotometrically the decolorization upon reduction of the co-substrate 2,6-dichlorophenolindophenol (DCIP), at 520 nm (ε_520_=6,800 M^-1^.cm^-1^). Most experiments were carried out at the optimal pH value of 5.5. Substrate specificity was assessed by screening 14 different substrates. Unless stated otherwise, reactions (100 μL final reaction volume) were carried out in 96-wells transparent microtiter plates (Corning Costar, Corning, NY, USA) and contained *Um*AA3_2-A (110 nM) and DCIP (0.4 mM) in citrate-phosphate (50 mM, pH 5.5). The mixtures were incubated during 2 min at 30 °C before the reaction was initiated by the addition of substrate (250 mM final for glucose and 2.5 mM for other substrates, including glucose). The absorbance was monitored over 10 min using a Tecan Infinite M200 (Tecan, Switzerland) plate reader. All reactions were carried out in triplicate. Initial rates, determined at various substrate concentrations, were used to calculate the kinetic parameters according to the standard Michaelis-Menten equation for G3G or using a modified model accounting for excess-substrate inhibition in the case of glucose. SigmaPlot 12.0 was used to fit the experimental data.

### Glycoside hydrolase activity assay

The activity of *Um*GH16_1-A was evaluated by monitoring the release of gluco-oligosaccharides from various glucans by high-performance anion exchange chromatography (HPAEC) coupled to pulsed amperometric detection (PAD) (*vide infra*). Unless stated otherwise, reactions (500 μL final reaction volume) were carried out in 2 mL Eppendorf tubes and contained the substrate (10 g.L^-1^) in citrate-phosphate buffer (50 mM, pH 5.5). The mixtures were incubated during 2 min at 30 °C in a Thermomixer (1,000 rpm) and the reactions were initiated by the addition of *Um*GH16_1-A_cd (10 nM). For each time point (15 min to 4 h), one sample (500 μL) is sacrificed by boiling for 10 min, centrifuged (12,000 *g*, 2 min, 4 °C), and diluted 10-fold in milliQ H2O before injection on the HPAEC column. Reactions using FCW extract were incubated overnight (16 to 18 h) and the supernatant was injected without prior dilution.

Reactions combining *Um*GH16_1-A_cd and *Um*AA3_2-A were carried out under similar conditions as described above with the addition of *Um*AA3_2-A (1 μM) and DCIP (400 μM).

### HPAEC-PAD analyses

The detection of soluble oligosaccharides is performed using HPAEC-PAD (DIONEX ICS6000 system, Thermo Fisher Scientific, Waltham, MA, USA). The system is equipped with a CarboPac-PA1 guard column (2 x 50 mm) and a CarboPac-PA1 column (2 x 250 mm) kept at 30 °C. Elution was carried out at a flow rate of 0.25 mL.min^-1^ and 25 μL of sample was injected. The eluents used were 100 mM NaOH (eluent A) and NaAc (1 M) in 100 mM NaOH (eluent B). The initial conditions were set to 100% eluent A, and the following gradient was applied: 0-10 min, 0-10% B; 10-35 min, 10-35% B (linear gradient); 35-40 min, 30-100% B (curve 6); 40-41 min, 100-0% B; 41-50 min, 100% A. Integration was performed using the Chromeleon 7.2.10 software based on commercially-available standards: laminari-oligosaccharides and G6G. G3G^ox^ and G6G^ox^ standards were respectively prepared by incubating G3G and G6G (1 mM each) with *Um*AA3_2-A (1 μM) and DCIP (2 mM) in citrate phosphate buffer (50 mM, pH 5.5), in a thermomixer (30 °C, 1,000 rpm) during 24 h.

### Linkage analyses

Polysaccharides (laminarin, pachyman and yeast β-glucans) were prepared at a concentration of 1 mg.mL^-1^ in dimethyl sulfoxide (DMSO) and left overnight at 60 °C under constant agitation. Methylation (method adapted from (51)) was performed with 500 μL of each sample by adding in the following order: 500 μL of NaOH-DMSO reagent and sonicate the tubes during 10 min, 100 μL of methyl iodide and sonicate the tubes during 10 min (twice) and 200 μL of methyl iodide and sonicate the tubes during 5 min. The reaction was stopped by the addition of H2O (2 mL) and the methylated products were extracted with chloroform (500 μL). The solutions were vigorously vortexed before a brief centrifugation, which allowed a strict separation of two phases. The aqueous supernatant phase was removed by aspiration. The organic phase was washed three times with H2O (2 mL). Methylated carbohydrates were hydrolyzed with 2 M trifluoroacetic acid in presence of an internal standard (myo-inositol) and converted to the corresponding alditol acetates. The partially methylated alditol acetates were analyzed by GC-MS (TRACE-GC-ISQ, Thermo^™^) on a non-polar thermo scientific™ TraceGOLD^™^ TG-1MS GC Column (30 m x 0.25 mm x 0.25 μm), carrier gas H2 at 1.5 mL.min^-1^. The sample was injected at 240 °C and the oven temperature was maintained for 5 min at 60 °C and increased up to 315 °C (3 °C /min), and further maintained at 315 °C for 2 min. The gas flow rate was set at 1.5 mL.min^-1^. The ion source temperature of the electron impact (EI) mass spectrometer was 230 °C. Masses were acquired with a scan range from m/z 100 to 500. Identification of partially methylated alditol acetates was based on their retention time and combined with confirmed by mass spectra fragmentation and compared to a home-made library. Quantitative detection was performed at 220 °C with a flame ionization detector (FID).

### Matrix assisted laser desorption/ionization (MALDI)-Time of flight (TOF) analysis

MALDI-TOF-MS spectra were acquired on a Rapiflex TissueTyper mass spectrometer (Bruker Daltonics, Bremen, Germany), equipped with a Smartbeam II Laser (355 nm, 10 kHz) and reflector detection. Samples were diluted in H2O (100 μg.mL^-1^) and directly mixed on a polished steel MALDI target plate with a solution of ionic liquid matrix DMA-DHB (2,5-dihydroxybenzoic acid 100 mg.mL^-1^ in H2O/ACN (50:50 vol/vol) with an addition of 0.2% of *N,N*-dimethylaniline (52). Spectra were recorded in the *m/z* range 350-3200 using FlexControl and processed using FlexAnalysis (Bruker Daltonics, Billerica, MA, USA). Mass spectra were acquired in positive ionization mode.

### Ultra High-Performance Liquid Chromatography (UHPLC)-Electrospray (ESI)-Ion trap (IT) analysis

UHPLC-ESI-IT acquisitions were performed on an amaZon SL 3D ion trap mass spectrometer (Bruker Daltonics, Bremen, Germany) coupled with an Acquity H-Class UHPLC (Waters, Wilmslow, UK). Samples were diluted in a solution of H2O/ACN (95.5:4.5) at 10 μg.mL^-1^. 10 μL of each sample was injected on an Hypercarb column (100 × 1 mm, particle size 3 μm, Thermo-Fisher Scientific, Courtaboeuf, France) heated at 80 °C with a flow rate settled at 0.165 mL.min^-1^. A binary gradient was performed. The gradient started with 8 min at 95.5% of A (H2O) and then ramped linearly to 80% of B (ACN) in 22 min and stayed at 80% of B during 12 min; initial conditions were restored during the last 5 min. The ESI source parameters were the following: capillary voltage: 4.5 kV; nebulizer gas: 7.3 psi; and dry gas: 4 L.min^-1^ (80 °C). Mass spectra were recorded in the *m/z* range 350-2200 in the positive ionization mode. Acquisitions were performed using TrapControl 8.0 and Compass HyStar 4.1 (Bruker Daltonics, Bremen, Germany). Data were processed using Data Analysis 4.4 (Bruker Daltonics, Bremen, Germany).

## ACKNOWLEDGEMENTS

We would like to thank Caroline Olivé for technical support during *Um*AA3_2-A cloning work and Elodie Drula (BBF/AFMB) for re-annotating the secretome of *U. maydis* and the CAZy team (AFMB) for providing us with the set of GH16 sequences underlying the phylogenetic tree. We thank Megazyme (now NEOGEN) for providing information on their *endo*-1,3-β-d-glucanase (E-LAMSE). We also thank the CIRM-CF for providing the *U. maydis* strain. MS analyses were performed on the BIBS instrumental platform (http://www.bibs.inrae.fr/bibs_eng/, UR1268 BIA, IBiSA, Phenome-Emphasis-FR ANR-11-INBS-0012, PROBE/CALIS infrastructures, Biogenouest). This study was supported by INRAE through the EvoFun project (PAF_02), the French National Agency for Research (“Agence Nationale de la Recherche”) through the “Projet de Recherche Collaboratif International” ANR-NSERC (FUNTASTIC project, ANR-17-CE07-0047). Part of the work described was performed using services provided by the 3PE platform, a member of IBISBA-FR (https://doi.org/10.15454/08BX-VJ91; www.ibisba.fr), the French node of the European research infrastructure, EU-IBISBA (www.ibisba.eu).

## AUTHOR CONTRIBUTIONS

J.-L.R., M.H., O.T. and S.G. carried out the enzymology experiments. D.R. and S.L.G. carried out mass spectrometry analyses. J.-L.R., D.R., S.L.G., G.S, J.-G.B. and B.B. interpreted the data. J.-G.B. and B.B. conceptualized the study, designed the experiments, and supervised the work. B.B. wrote the first draft and finalized the manuscript. All authors contributed to the writing of the manuscript, with main contributions from J.-L.R. and J.-G.B. All authors reviewed and approved the final version of the manuscript.

## REFERENCES

1. Lebreton A, Zeng Q, Miyauchi S, Kohler A, Dai YC, Martin FM. 2021. Evolution of the Mode of Nutrition in Symbiotic and Saprotrophic Fungi in Forest Ecosystems. Annu Rev Ecol Evol Syst. Annual Reviews 52:385–404.

2. Grigoriev I V., Nikitin R, Haridas S, Kuo A, Ohm R, Otillar R, Riley R, Salamov A, Zhao X, Korzeniewski F, Smirnova T, Nordberg H, Dubchak I, Shabalov I. 2014. MycoCosm portal: Gearing up for 1000 fungal genomes. Nucleic Acids Res 42:D699–D704.

3. Martinez D, Challacombe J, Morgenstern I, Hibbett D, Schmoll M, Kubicek CP, Ferreira P, Ruiz-Duenas FJ, Martinez AT, Kersten P, Hammel KE, Vanden Wymelenberg A, Gaskell J, Lindquist E, Sabat G, Splinter BonDurant S, Larrondo LF, Canessa P, Vicuna R, Yadav J, Doddapaneni H, Subramanian V, Pisabarro AG, Lavín JL, Oguiza JA, Master E, Henrissat B, Coutinho PM, Harris P, Magnuson JK, Baker SE, Bruno K, Kenealy W, Hoegger PJ, Kües U, Ramaiya P, Lucas S, Salamov A, Shapiro H, Tu H, Chee CL, Misra M, Xie G, Teter S, Yaver D, James T, Mokrejs M, Pospisek M, Grigoriev I V, Brettin T, Rokhsar D, Berka R, Cullen D. 2009. Genome, transcriptome, and secretome analysis of wood decay fungus Postia placenta supports unique mechanisms of lignocellulose conversion. Proc Natl Acad Sci 106:1954–1959.

4. Vanden Wymelenberg A, Gaskell J, Mozuch M, Kersten P, Sabat G, Martinez D, Cullen D. 2009. Transcriptome and secretome analyses of Phanerochaete chrysosporium reveal complex patterns of gene expression. Appl Environ Microbiol 75:4058–4068.

5. Eastwood DC, Floudas D, Binder M, Majcherczyk A, Schneider P, Aerts A, Asiegbu FO, Baker SE, Barry K, Bendiksby M, Blumentritt M, Coutinho PM, Cullen D, de Vries RP, Gathman A, Goodell B, Henrissat B, Ihrmark K, Kauserud H, Kohler A, LaButti K, Lapidus A, Lavin JL, Lee Y-H, Lindquist E, Lilly W, Lucas S, Morin E, Murat C, Oguiza JA, Park J, Pisabarro AG, Riley R, Rosling A, Salamov A, Schmidt O, Schmutz J, Skrede I, Stenlid J, Wiebenga A, Xie X, Kües U, Hibbett DS, Hoffmeister D, Högberg N, Martin F, Grigoriev I V., Watkinson SC. 2011. The plant cell wall-decomposing machinery underlies the functional diversity of forest fungi. Science 333:762–765.

6. Rytioja J, Hildén K, Yuzon J, Hatakka A, de Vries RP, Mäkelä MR. 2014. Plant-polysaccharide-degrading enzymes from Basidiomycetes. Microbiol Mol Biol Rev 78:614–649.

7. Bissaro B, Varnai A, Røhr ÅK, Eijsink VGH. 2018. Oxidoreductases and reactive oxygen species in lignocellulose biomass conversion. Microbiol Mol Biol Rev 82:e00029–18.

8. Hage H, Miyauchi S, Virágh M, Drula E, Min B, Chaduli D, Navarro D, Favel A, Norest M, Lesage-Meessen L, Bálint B, Merényi Z, de Eugenio L, Morin E, Martínez A, Baldrian P, Štursová M, Martínez MJ, Novotny C, Magnuson J, Spatafora J, Maurice S, Pangilinan RM. 2021. Gene family expansions and transcriptome signatures uncover fungal adaptations to wood decay. Enviromental Microbiol 23:5716–5732.

9. Miyauchi S, Navarro D, Grisel S, Chevret D, Berrin JG, Rosso MN. 2017. The integrative omics of white-rot fungus Pycnoporus coccineus reveals co-regulated CAZymes for orchestrated lignocellulose breakdown. PLoS One 12:e0175528.

10. Couturier M, Navarro D, Olivé C, Chevret D, Haon M, Favel A, Lesage-Meessen L, Henrissat B, Coutinho PM, Berrin JG. 2012. Post-genomic analyses of fungal lignocellulosic biomass degradation reveal the unexpected potential of the plant pathogen Ustilago maydis. BMC Genomics 13:57.

11. Roncero C, Vázquez de Aldana CR. 2019. Glucanases and Chitinases. Curr Top Microbiol Immunol 425:131–166.

12. Emri T, Molnár Z, Szilágyi M, Pócsi I. 2008. Regulation of autolysis in Aspergillus nidulans. Appl Biochem Biotechnol 151:211–220.

13. Gow NAR, Latge J-P, Munro CA. 2017. The Fungal Cell Wall: Structure, Biosynthesis, and Function. Microbiol Spectr 5.

14. Ruiz-Herrera J, Ortiz-Castellanos L. 2019. Cell wall glucans of fungi. A review. Cell Surf 5:100022.

15. Dumas-Gaudot E, Slezack S, Dassi B, Pozo MJ, Gianinazzi-Pearson V, Gianinazzi S. 1996. Plant hydrolytic enzymes (chitinases and β-1,3-glucanases) in root reactions to pathogenic and symbiotic microorganisms, p. 211–221. In Plant and Soil.

16. Dean R, Van Kan JAL, Pretorius ZA, Hammond-Kosack KE, Di Pietro A, Spanu PD, Rudd JJ, Dickman M, Kahmann R, Ellis J, Foster GD. 2012. The Top 10 fungal pathogens in molecular plant pathology. Mol Plant Pathol 13:414–430.

17. Kämper J, Kahmann R, Bölker M, Ma LJ, Brefort T, Saville BJ, Banuett F, Kronstad JW, Gold SE, Müller O, Perlin MH, Wösten HAB, De Vries R, Ruiz-Herrera J, Reynaga-Peña CG, Snetselaar K, McCann M, Pérez-Martín J, Feldbrügge M, Basse CW, Steinberg G, Ibeas JI, Holloman W, Guzman P, Farman M, Stajich JE, Sentandreu R, González-Prieto JM, Kennell JC, Molina L, Schirawski J, Mendoza-Mendoza A, Greilinger D, Münch K, Rössel N, Scherer M, Vran?s M, Ladendorf O, Vincon V, Fuchs U, Sandrock B, Meng S, Ho ECH, Cahill MJ, Boyce KJ, Klose J, Klosterman SJ, Deelstra HJ, Ortiz-Castellanos L, Li W, Sanchez-Alonso P, Schreier PH, Häuser-Hahn I, Vaupel M, Koopmann E, Friedrich G, Voss H, Schlüter T, Margolis J, Platt D, Swimmer C, Gnirke A, Chen F, Vysotskaia V, Mannhaupt G, Güldener U, Münsterkötter M, Haase D, Oesterheld M, Mewes HW, Mauceli EW, DeCaprio D, Wade CM, Butler J, Young S, Jaffe DB, Calvo S, Nusbaum C, Galagan J, Birren BW. 2006. Insights from the genome of the biotrophic fungal plant pathogen Ustilago maydis. Nature 444:97–101.

18. Levasseur A, Drula E, Lombard V, Coutinho PM, Henrissat B. 2013. Expansion of the enzymatic repertoire of the CAZy database to integrate auxiliary redox enzymes. Biotechnol Biofuels 6:41.

19. Viborg AH, Terrapon N, Lombard V, Michel G, Czjzek M, Henrissat B, Brumer H. 2019. A subfamily roadmap of the evolutionarily diverse glycoside hydrolase family 16 (GH16). J Biol Chem 294:15973–15986.

20. Drula E, Garron ML, Dogan S, Lombard V, Henrissat B, Terrapon N. 2022. The carbohydrate-active enzyme database: Functions and literature. Nucleic Acids Res 50:D571–D577.

21. Rizzi YS, Happel P, Lenz S, Urs MJ, Bonin M, Cord-Landwehr S, Singh R, Moerschbacher BM, Kahmann R. 2021. Chitosan and chitin deacetylase activity are necessary for development and virulence of ustilago maydis. MBio 12:1–18.

22. Lanver D, Müller AN, Happel P, Schweizer G, Haas FB, Franitza M, Pellegrin C, Reissmann S, Altmüller J, Rensing SA, Kahmann R. 2018. The biotrophic development of ustilago maydis studied by RNA-seq analysis. Plant Cell 30:300–323.

23. Bakke M, Kamei JI, Obata A. 2011. Identification, characterization, and molecular cloning of a novel hyaluronidase, a member of glycosyl hydrolase family 16, from Penicillium spp. FEBS Lett 585:115–120.

24. Qin Z, Yang S, Zhao L, You X, Yan Q, Jiang Z. 2017. Catalytic mechanism of a novel glycoside hydrolase family 16 “elongating” β-transglycosylase. J Biol Chem 292:1666–1678.

25. Sützl L, Laurent CVFP, Abrera AT, Schütz G, Ludwig R, Haltrich D. 2018. Multiplicity of enzymatic functions in the CAZy AA3 family. Appl Microbiol Biotechnol 102:2477–2492.

26. Piumi F, Levasseur A, Navarro D, Zhou S, Mathieu Y, Ropartz D, Ludwig R, Faulds CB, Record E, 2014. A novel glucose dehydrogenase from the white-rot fungus Pycnoporus cinnabarinus: production in Aspergillus niger and physicochemical characterization of the recombinant enzyme. Appl Microbiol Biotechnol 98:10105–10118.

27. Cerutti G, Gugole E, Montemiglio LC, Turbé-Doan A, Chena D, Navarro D, Lomascolo A, Piumi F, Exertier C, Freda I, Vallone B, Record E, Savino C, Sciara G. 2021. Crystal structure and functional characterization of an oligosaccharide dehydrogenase from Pycnoporus cinnabarinus provides insights into fungal breakdown of lignocellulose. Biotechnol Biofuels 2021 141 14:1–18.

28. Jumper J, Evans R, Pritzel A, Green T, Figurnov M, Ronneberger O, Tunyasuvunakool K, Bates R, Žídek A, Potapenko A, Bridgland A, Meyer C, Kohl SAA, Ballard AJ, Cowie A, Romera-Paredes B, Nikolov S, Jain R, Adler J, Back T, Petersen S, Reiman D, Clancy E, Zielinski M, Steinegger M, Pacholska M, Berghammer T, Bodenstein S, Silver D, Vinyals O, Senior AW, Kavukcuoglu K, Kohli P, Hassabis D. 2021. Highly accurate protein structure prediction with AlphaFold. Nature 596:583–589.

29. Vasur J, Kawai R, Andersson E, Igarashi K, Sandgren M, Samejima M, Ståhlberg J. 2009. X-ray crystal structures of Phanerochaete chrysosporium Laminarinase 16A in complex with products from lichenin and laminarin hydrolysis. FEBS J 276:3858–3869.

30. Kawai R, Igarashi K, Yoshida M, Kitaoka M, Samejima M. 2006. Hydrolysis of β-1,3/1,6-glucan by glycoside hydrolase family 16 endo-1,3(4)-β-glucanase from the basidiomycete Phanerochaete chrysosporium. Appl Microbiol Biotechnol 71:898–906.

31. Dita Wijayanti S, Sützl L, Duval A, Haltrich D, Phylogenetic Clades UJ, Bankole O, Schlosser D. 2021. Characterization of Fungal FAD-Dependent AA3_2 Glucose Oxidoreductases from Hitherto Unexplored Phylogenetic Clades. J Fungi 2021, Vol 7, Page 873 7:873.

32. Igarashi K, Samejima M, Eriksson KEL. 1998. Cellobiose dehydrogenase enhances Phanerochaete chrysosporium cellobiohydrolase I activity by relieving product inhibition. Eur J Biochem 253:101–106.

33. Kracher D, Scheiblbrandner S, Felice AKG, Breslmayr E, Preims M, Haltrich D, Eijsink VGH, Ludwig R. 2016. Extracellular electron transfer systems fuel cellulose oxidative degradation. Science 352:1098–1101.

34. Haddad Momeni M, Fredslund F, Bissaro B, Raji O, Vuong T V., Meier S, Nielsen TS, Lombard V, Guigliarelli B, Biaso F, Haon M, Grisel S, Henrissat B, Welner DH, Master ER, Berrin JG, Abou Hachem M. 2021. Discovery of fungal oligosaccharide-oxidising flavo-enzymes with previously unknown substrates, redox-activity profiles and interplay with LPMOs. Nat Commun 12.

35. Garajova S, Mathieu Y, Beccia MR, Bennati-Granier C, Biaso F, Fanuel M, Ropartz D, Guigliarelli B, Record E, Rogniaux H, Henrissat B, Berrin J-G. 2016. Single-domain flavoenzymes trigger lytic polysaccharide monooxygenases for oxidative degradation of cellulose. Sci Rep 6:1–9.

36. Buckeridge MS, Rayon C, Urbanowicz B, Tiné MAS, Carpita NC. 2004. Mixed Linkage (1→3),(1→4)-β-d-Glucans of Grasses. Cereal Chem 81:115–127.

37. Zhang Q, Cheetamun R, Dhugga KS, Rafalski JA, Tingey S V., Shirley NJ, Taylor J, Hayes K, Beatty M, Bacic A, Burton RA, Fincher GB. 2014. Spatial gradients in cell wall composition and transcriptional profiles along elongating maize internodes. BMC Plant Biol 14:1–19.

38. Novak M, Vetvicka V. 2008. β-glucans, history, and the present: Immunomodulatory aspects and mechanisms of action. J Immunotoxicol. Taylor & Francis https://doi.org/10.1080/15476910802019045.

39. Trouvelot S, Héloir MC, Poinssot B, Gauthier A, Paris F, Guillier C, Combier M, Trdá L, Daire X, Adrian M. 2014. Carbohydrates in plant immunity and plant protection: Roles and potential application as foliar sprays. Front Plant Sci 5:592.

40. Wagener J, Striegler K, Wagener N. 2020. α-and β-1,3-Glucan Synthesis and Remodeling. Curr Top Microbiol Immunol 425:53–82.

41. Vandhana TM, Reyre J-L, Dangudubiyyam S, Berrin J-G, Bissaro B, Madhuprakash J. 2022. On the expansion of biological functions of Lytic Polysaccharides Monooxygenases. New Phytol 233:2380–2396.

42. Sabbadin F, Urresti S, Henrissat B, Avrova AO, Welsh LRJ, Lindley PJ, Csukai M, Squires JN, Walton PH, Davies GJ, Bruce NC, Whisson SC, McQueen-Mason SJ. 2021. Secreted pectin monooxygenases drive plant infection by pathogenic oomycetes. Science 373:774–779.

43. Kretschmer M, Damoo D, Sun S, Lee CWJ, Croll D, Brumer H, Kronstad J. 2022. Organic acids and glucose prime late-stage fungal biotrophy in maize. Science 376:1187–1191.

44. Haon M, Grisel S, Navarro D, Gruet A, Berrin JG, Bignon C. 2015. Recombinant protein production facility for fungal biomass-degrading enzymes using the yeast *Pichia pastoris*. Front Microbiol 6:1–12.

45. Couturier M, Haon M, Coutinho PM, Henrissat B, Lesage-Meessen L, Berrin JG. 2011. Podospora anserina hemicellulases potentiate the Trichoderma reesei secretome for saccharification of lignocellulosic biomass. Appl Environ Microbiol 77:237–246.

46. Hall TA. 1999. BIOEDIT: a user-friendly biological sequence alignment editor and analysis program for Windows 95/98/ NT. Nucleic Acids Symp Ser 41:95–98.

47. Katoh K, Rozewicki J, Yamada KD. 2018. MAFFT online service: Multiple sequence alignment, interactive sequence choice and visualization. Brief Bioinform 20:1160–1166.

48. Whelan S, Goldman N. 2001. A general empirical model of protein evolution derived from multiple protein families using a maximum-likelihood approach. Mol Biol Evol 18:691–699.

49. Letunic I, Bork P. 2019. Interactive Tree of Life (iTOL) v4: Recent updates and new developments. Nucleic Acids Res 47:256–259.

50. Navarro D, Chaduli D, Taussac S, Lesage-Messeen L, Grisel S, Haon M, Callac P, Courtecuisse R, Decock C, Dupont J, Richard-Forget F, Fournier J, Guinberteau J, Lechat C, Moreau P-A, Pinson-Gadais L, Rivoire B, Sage L, Welti S, Rosso M-N, Berrin J-G, Bissaro B, Favel A. 2021. Large-scale phenotyping of 1,000 fungal strains for the degradation of non-natural, industrial compounds. Commun Biol 4:871.

51. Pettolino FA, Walsh C, Fincher GB, Bacic A. 2012. Determining the polysaccharide composition of plant cell walls. Nat Protoc 7:1590–1607.

52. Ropartz D, Bodet PE, Przybylski C, Gonnet F, Daniel R, Fer M, Helbert W, Bertrand D, Rogniaux H. 2011. Performance evaluation on a wide set of matrix-assisted laser desorption ionization matrices for the detection of oligosaccharides in a high-throughput mass spectrometric screening of carbohydrate depolymerizing enzymes. Rapid Commun Mass Spectrom 25:2059–2070.

